# Cutting through the clutter: minimizing redundancy in GO enrichment analysis with evoGO

**DOI:** 10.1101/2025.02.24.639258

**Authors:** Ievgen Strielkov, Greta Buinovskaja, Alexey Uvarovskii, Vladimir Galatenko

## Abstract

Gene Ontology (GO) enrichment analysis is a powerful tool for elucidating underlying biological processes in high-throughput transcriptomic and proteomic studies. However, the redundancy of identified enriched GO terms, caused by the hierarchical nature of GO, significantly complicates the prioritization of relevant terms and makes drawing concise conclusions challenging. To address this problem, we developed evoGO, a novel method that aims to improve the specificity and relevance of enrichment results by considering the GO hierarchy during the analysis. Built on the foundation of conventional overrepresentation analysis (ORA), evoGO reduces the impact of differentially expressed genes on the significance of a given GO term if those genes already contribute to a higher significance of any descendant GO term. The effectiveness of the algorithm was evaluated against other advanced ORA-based GO enrichment analysis methods (topGO, clusterProfiler, and SetRank) using synthetic and real-life datasets. In the synthetic benchmarks, evoGO reduced the number of enriched terms identified by ORA on average by 30% while recovering the highest fraction (96%) of the true positives. When applied to real-life data, evoGO most efficiently prioritized tissue-specific GO terms in the analyses aimed to capture the biological processes inherent to various human tissues. Furthermore, evoGO eliminated the fewest enriched GO terms that had no significantly enriched relatives and thus should not be considered redundant. Finally, evoGO was one of the fastest of the tested methods and, unlike competitors, maintained consistent execution times across both benchmarks. In conclusion, our findings demonstrate that among tested ORA-based approaches, evoGO stands out as an effective and fast method for minimizing redundancy in GO enrichment analysis results while reliably preserving biologically relevant information. We believe that evoGO could be a valuable tool for conducting downstream analyses, particularly in the context of high-throughput transcriptomic and proteomic screening studies. The evoGO method has been implemented as an R package, which is available on GitHub.

## INTRODUCTION

Genome-wide gene expression studies are commonly used to investigate the molecular mechanisms underlying a phenotype of interest. These studies are known to produce vast amounts of data and depend on specific analysis techniques. In particular, given the large number of known genes, grouping them into sets by common features, such as function, related biological process, or cell structure, facilitates the interpretation of results. Specifically, identifying gene sets enriched with genes of interest (e.g., differentially expressed genes) can help pinpoint the key biological differences between two experimental conditions.

A number of databases currently provide extensive, curated collections of functional gene sets [1, 2, 3]. Among them, Gene Ontology (GO) is known to be one of the largest and most popular. Currently, it contains annotations for 1.5 million gene products of more than 5 thousand species [4]. GO consists of three independent domains: “biological process”, “molecular function”, and “cellular component”. Each domain is represented by a directed acyclic graph, with nodes corresponding to GO terms and edges indicating relationships between them. This forms a hierarchical structure where each term can have one or more parent terms.

Two classic enrichment analysis methods are commonly used with GO gene sets: overrepresentation analysis (ORA) and gene set enrichment analysis (GSEA). The first method tests whether a given gene set contains a disproportionately high number of differentially expressed (DE) genes using Fisher’s exact test (FET) or the chi-squared test [5]. The main disadvantage of this approach is that it relies on the DE/non-DE gene dichotomy, and thus its results strongly depend on the chosen significance threshold. In contrast, GSEA circumvents this problem by testing whether genes of a certain set tend to have higher (or lower) significance scores compared to the rest of the genes. Nevertheless, recent comparative studies have shown that ORA outperforms GSEA or similar functional class scoring methods when applied to real-life data [6, 7, 8, 9]. These findings, along with the simplicity of usage and ease of result interpretation, may explain the longstanding popularity of ORA.

Despite ORA’s widespread usage, this method, as well as GSEA, has several methodological shortcomings when applied to GO [discussed in 7, 10]. Arguably, one of the most important issues is the significant redundancy in the resulting lists of enriched terms [10, 11, 12]. The redundancy is caused by numerous overlaps in gene sets and is a common problem for pathway databases, including KEGG and Reactome [10]. However, it is most prominent for GO due to the complex hierarchical structure of its domains and the rule that genes attributed to a child term are also implicitly attributed to all parent terms.

Such ontology organization creates iterative gene set overlaps, which sometimes cause dozens of closely related terms to be enriched at the same time [10, 11, 12]. This significantly complicates the prioritization of relevant GO terms, making it more difficult to draw concise conclusions from the analysis results. Note that simply sorting gene sets by their p-values is not helpful in this situation, as more general GO terms often have higher significance than more specific ones. This is a direct consequence of the “gene inheritance” principle mentioned above.

A variety of enrichment analysis methods have been suggested to minimize redundancy in enrichment analysis results. These methods can generally be categorized into three groups. First, there are methods that rely on pre-simplified GO or modified gene sets [12, 13, 14, 15]. A potential disadvantage of this approach is that modifications to the original gene sets may result in a loss of certain valuable insights from the analysis, depending on the chosen simplification method [discussed in 16]. Other methods perform clustering of gene sets using various similarity metrics. This allows the identification of distinct “families” of enriched GO terms, which facilitates result interpretation [11, 17, 18]. However, such methods are typically not designed to improve the prioritization of terms within clusters.

Finally, some methods attempt to reduce the significance of redundant gene sets during the analysis [10, 19, 20]. Although the definition of a redundant term varies among different methods, this approach appears most promising, as it aims to improve term prioritization while preserving the original gene sets.

Following this idea, methods such as PADOG or SetRank down-weight genes that are attributed to multiple sets [10, 20]. When applied to GO, these methods treat terms as independent gene sets and ignore their hierarchy. Despite certain improvements reported by the authors for a non-hierarchical database (KEGG), decreasing the weights of common genes in unrelated gene sets can lead to biased results. Genes that are involved in multiple biological processes may represent key regulators, and down-weighting them can obscure their functional significance. Furthermore, reducing the role of certain genes can make the analysis more susceptible to noise and variability coming from the remaining DE genes.

The authors of the topGO method adopt a different approach. To our knowledge, topGO is currently the only method designed to deprioritize redundant enriched terms while considering the GO hierarchy [19]. The “weight” algorithm of the topGO R package adjusts the weights of DE genes between neighboring GO terms in a way that reinforces the differences in term significance. While we believe this is a sensible approach, topGO may have a few drawbacks. The iterative reciprocal up- and down-weighting of genes attributed to neighboring terms performed by “weight” not only complicates the interpretation of results, but may also unintentionally amplify random effects, such as noise, thereby increasing the proportion of false positives. Furthermore, the lengthy execution times of topGO analysis make it impractical or even unrealistic to use the package in high-throughput studies, which involve hundreds or thousands differential expression analyses, e.g., in drug screening.

Against this background, we present evoGO, a GO enrichment analysis method that maintains the power and interpretability of ORA, while aiming to efficiently reduce redundancy in results without compromising their validity. While implementing the algorithm as an R package, we made a special effort to minimize the analysis execution time. This enables evoGO usage for high-throughput RNA sequencing and proteomics studies. Additionally, one of our priorities was to ensure the reproducibility of analyses. The evoGO R package allows users to choose a specific GO release as well as the version of gene-to-term mapping. To validate our method, in this paper, we compare its performance to that of several popular versions of ORA using synthetic and real-life dataset.

## METHODS

### evoGO algorithm

The original ORA for a gene set is conducted using a 2×2 contingency table containing the number of DE genes and non-DE genes that are related and not related to the set. When applied to such a table, FET assesses the null hypothesis that the term and phenotype are independent. The evoGO algorithm builds on this approach by considering the GO term hierarchy in order to reduce the significance of potentially redundant, less specific terms.

The algorithm accepts a list of GO term relationships (only *“is a”* relationships are considered), gene-to-term mapping, a list of genes of interest (DE genes), and a background gene list as input. A full set of ancestors is identified, and the shortest distance to the root term of the current domain is calculated for each term. Next, genes are assigned to terms based on the provided annotation. According to the GO inheritance rule, all genes of child terms are also added to parent terms. This procedure is performed iteratively, starting from terms of the lowest level, i.e., the terms with the greatest distance to the root term. The initial weight for each gene is set to 1. The terms that have zero or only a few genes attributed to them (by default, fewer than 5) are removed, and their relations to other terms are ignored during the analysis. Due to gene inheritance rule, the removed GO terms are the most specific terms in the GO hierarchy. This step concludes data preparation. evoGO begins the analysis by evaluating the initial significance of each GO term independently using FET. This procedure is equivalent to standard ORA. Next, the algorithm iterates over GO terms, moving from the lowest level of the graph to its root. For each term, evoGO finds its less significant ancestors (all directly and indirectly related higher level GO terms) and calculates the corresponding down-weighting coefficients.

These coefficients represent the ratio of the current term’s p-value to the ancestor term’s p-value. The obtained coefficients are applied to the weights of those ancestor genes that are also associated with the current term. Changing the gene weights based on the ratio of GO term p-values was also previously suggested by the authors of the topGO algorithm [19]. After evoGO reaches the end of the current level, the updated weights are used to recalculate p-values for affected ancestors. For this, the sums of term gene weights are rounded to a whole number and used instead of DE gene counts in FET. The described procedure is performed for each level of the GO graph. In its output, evoGO provides both standard “unweighted” ORA p-values and the p-values obtained considering gene weights. Notably, unlike topGO, evoGO avoids iterative bidirectional weight adjustments between neighboring terms. This preserves the original p-value of the most significant term in a given GO ancestry, making the interpretation of the resulting p-like scores easier. In particular, conventional significance thresholds can still be applied similarly to ordinary ORA.

### Implementation and availability

The evoGO method has been implemented as an R package. The current version of the package uses the Ensembl database to annotate genes to GO terms. In future updates, we aim to expand the available options for gene annotation by incorporating other sources. To ensure the reproducibility of analyses, the evoGO package provides users with the ability to easily select specific releases of both the Gene Ontology and the Ensembl database, a feature often overlooked by similar tools. Additionally, evoGO allows users to utilize custom feature sets for GO term annotation. The evoGO package is publicly available on GitHub (https://github.com/Evotec-Bioinformatics/evoGO). Installation and usage instructions, as well as sample code, can be found within the README file in the repository.

### Other tested methods

In order to evaluate the performance of the evoGO method, we compared it to that of other popular methods of GO enrichment analysis. Considering that the aim of our work was to improve upon existing ORA, we included other ORA-based methods in our benchmark to ensure adequate comparability of results. Besides the basic FET, for our testing, we chose the following methods.

### topGO

The topGO R package was introduced in 2006 by *Alexa et al*. and remains one of the most popular R packages in the field [19]. It features several algorithms for GO enrichment analysis, among which “*weight*” was shown to be the most advanced. Similarly to evoGO, the “*weight*” algorithm assigns weights to genes based on the ratios of significance scores of neighboring GO terms. This approach aims to reinforce differences in significance between a node and its neighbors. If a node is more significant, genes in its children are down-weighted, and vice versa. The final significance score of a GO term is computed using a weighted contingency table.

### SetRank

The SetRank algorithm aims to reduce redundancy in gene set enrichment analysis by eliminating gene sets that appear significant solely due to their overlap with another significant gene set [10]. This method does not rely on a gene set hierarchy and is designed to work with any plain gene sets. For each set in a collection, a primary p-value is calculated using FET (if no gene ranks are provided). Then, gene sets that derive their significance primarily from their overlap with another gene set are discarded. This step aims to reduce false positives in the results. The remaining gene sets are presented as a directed graph. In this structure, each node corresponds to a gene set, and the edges indicate intersections between these sets. The direction of the edges indicates the relative significance of the intersecting gene sets. They are directed from the less significant to the more significant set. To prioritize the gene sets for further evaluation, a score called the SetRank value is calculated for each gene set in the network using the PageRank algorithm. Finally, p-values are calculated using the obtained SetRank values.

### clusterProfiler

The clusterProfiler package offers advanced tools for classical ORA and GSEA [21]. In particular, it offers functionality to minimize redundancy in GO enrichment analysis results with the “*simplify*” function. This function uses the GOSemSim package to calculate semantic similarities among enriched GO terms based on information content or graph structure. The function then removes GO terms with similarity values above a certain threshold, retaining only one representative term for each group of similar terms. The representative term can be selected based on the highest significant enrichment score or another criterion.

### Synthetic benchmark

As the first step in our comparison, we evaluated the ability of the tested GO enrichment analysis methods to detect enriched terms and eliminate redundant ones. To accomplish this, we performed analyses using synthetic data. We randomly selected 5-50 GO terms from the “biological process” domain with 10-300 genes attributed to them. We believe that terms containing more than 300 genes are too general to provide meaningful results. At the same time, these terms constitute only a very minor fraction of all GO terms in the domain. From each selected term, we marked 10%, 20%, and 30% of genes as differentially expressed. Typically, this resulted in 30 - 800 selected genes, depending on the number of sampled GO terms. We repeated this procedure 500 times. For each set of selected genes, we performed GO enrichment analysis using the methods described above. In the obtained results, we evaluated the fraction of recovered preselected GO terms, fractions of their recovered ancestor and descendant terms, as well as the fraction of recovered unrelated terms.

A potential flaw of the described benchmark is that other terms that appear enriched alongside the preselected (“true positive”) terms may not necessarily be “false positives,” as the similarity between their gene sets may represent the actual biological truth.

However, we believe that this procedure enables the comparison of the general sensitivity and selectivity of the tested methods. For example, certain methods may consistently return more highly enriched child or parent terms, fail to recover the key enriched terms, or identify completely irrelevant GO terms as enriched.

While analyzing the results obtained using the described method, we considered non-adjusted significance scores for the identification of enriched terms. We believe that there are several reasons why using non-adjusted scores is a more suitable approach for our purposes.

#### Interdependence of GO Terms

Traditional p-value correction methods, such as Benjamini-Hochberg correction, assume independence of each statistical test performed. However, the hierarchical structure of the GO and the genes shared among the GO terms make the tests interdependent. In this situation, p-value adjustment methods can be overly conservative, reducing the power of the test.

#### Number of Returned Terms

Different GO enrichment methods filter out certain GO terms at different stages of the analysis. Consequently, the tested method may perform a significantly different number of statistical tests, potentially leading to a biased effect of p-value adjustment, such as over-correction for methods that return more terms.

#### P-Value-Like Scores

Most of the tested methods do not return true p-values. Instead, they produce p-value-like scores by manipulating original p-values. Classical p-value correction methods are designed to control error rates based on the assumption that p-values represent the probability of observing the data (or more extreme data) under the null hypothesis. However, as some of the p-values could be artificially changed (e.g., increased for redundant GO terms) by the methods, the assumption that underlies these correction methods no longer holds. Therefore, the error rates that these corrections aim to control may not be accurately controlled, which could lead to misleading results.

Considering all the reasons mentioned above, we used the original significance scores (p) returned by the tested methods in our analysis. A GO term was considered significantly enriched if p was less than 0.05. Additionally, we compared the results obtained with different *p*value thresholds to ensure the consistency of the outcomes.

In general, we recommend applying the Benjamini-Hochberg correction to the p-values returned by evoGO for routine analyses. Our observations suggest that this approach reduces the number of redundant enriched GO terms and decreases the fraction of false-positive terms recovered.

### Real data benchmark

Benchmarking GO enrichment analysis methods using real-life biological data often poses a significant challenge, primarily due to the lack of a fully established “ground truth”. The underlying mechanisms of a specific phenotype or disease are often not completely understood, making it difficult to evaluate the accuracy and reliability of a given method.

Moreover, biological processes are frequently complex and multifaceted, allowing for multiple plausible hypotheses or interpretations. This complicates discerning whether a particular method reflects the actual biological truth or just one of several possible perspectives.

Considering these challenges, we attempted to design a benchmarking procedure that relies solely on well-established biological knowledge. Specifically, we assumed that our current understanding of normal tissue-specific (TS) biological processes provides a more robust foundation for benchmarking than disease-associated experimental data, which is typically used for this purpose. In this context, we utilized the fact that the “biological process” domain of the Gene Ontology contains a significant fraction of TS terms.

Following this idea, we performed GO enrichment analyses using DE genes identified in comparisons of gene expression profiles of different human tissues. Finally, we compared the sets of recovered “true positive” TS GO terms provided by the tested methods.

### Gene expression in human tissues

The human tissue gene expression data used in this study were obtained from the Genotype-Tissue Expression (GTEx) database, release V8 (https://www.gtexportal.org/). GTEx is a comprehensive public database that catalogues gene expression across a wide range of human tissues [22]. For the primary benchmark, we used samples from six healthy human tissues. When selecting tissue samples from the database, we only considered those with a tissue degradation level of 0 or 1 and causes of death corresponding to 1 or 2 on the Hardy scale. In total, 29 heart, 19 brain, 14 lung, 25 skeletal muscle, 26 thyroid tissue, and 9 testis samples were selected for the benchmark.

### Differential expression analysis

The differential expression analysis was conducted using the DESeq2 package (version 1.38.2) in the R programming language (version 4.2.0) [23]. P-values were adjusted for multiple testing using the Benjamini-Hochberg (FDR) procedure. Genes were considered differentially expressed if their FDR-adjusted p-values were less than 0.05. For the GO enrichment analyses, we used a variable number of the top DE genes (100 - 3000) to compare method performance under different conditions.

### Tissue-specific GO terms

The following GO terms, each with at least 5 genes attributed to them, were considered TS: terms containing “heart”, “cardio”, and “cardiac” in their names were categorized as heart-specific terms (518 terms); terms containing “brain”, “neuro”, and “cerebral” were categorized as brain-specific terms (466 terms); terms containing “lung”, “pneumo”, and “pulmonary” were categorized as lung-specific terms (90 terms). Terms containing keywords attributed to multiple tissues were excluded. Although some of the keywords may not be exclusively associated with the given tissue from a biological perspective, we believe that, considering the fundamental functional and developmental differences between the brain, heart, and lung, the selected terms could be attributed to only one of the three tissues in our benchmark. For example, even though the “neuro” keyword in a GO term may not be uniquely linked to the brain, but also to other parts of the nervous system, in the given context, it is unlikely to be truly enriched in a comparison between gene expression profiles of heart and lung tissues.

## RESULTS AND DISCUSSION

### Synthetic benchmark

The synthetic benchmark results revealed significant differences in the performance of the tested methods. As expected, FET yielded the highest number of enriched GO terms in all scenarios. Furthermore, this method recovered the highest fraction of true positive (TP) terms, i.e. the terms that had DE genes assigned to them at the initial phase of the benchmarking procedure (Fig. 1). In particular, FET identified practically all TPs when the fraction of DE genes in them was 0.2 or 0.3.

**Fig. 1.**
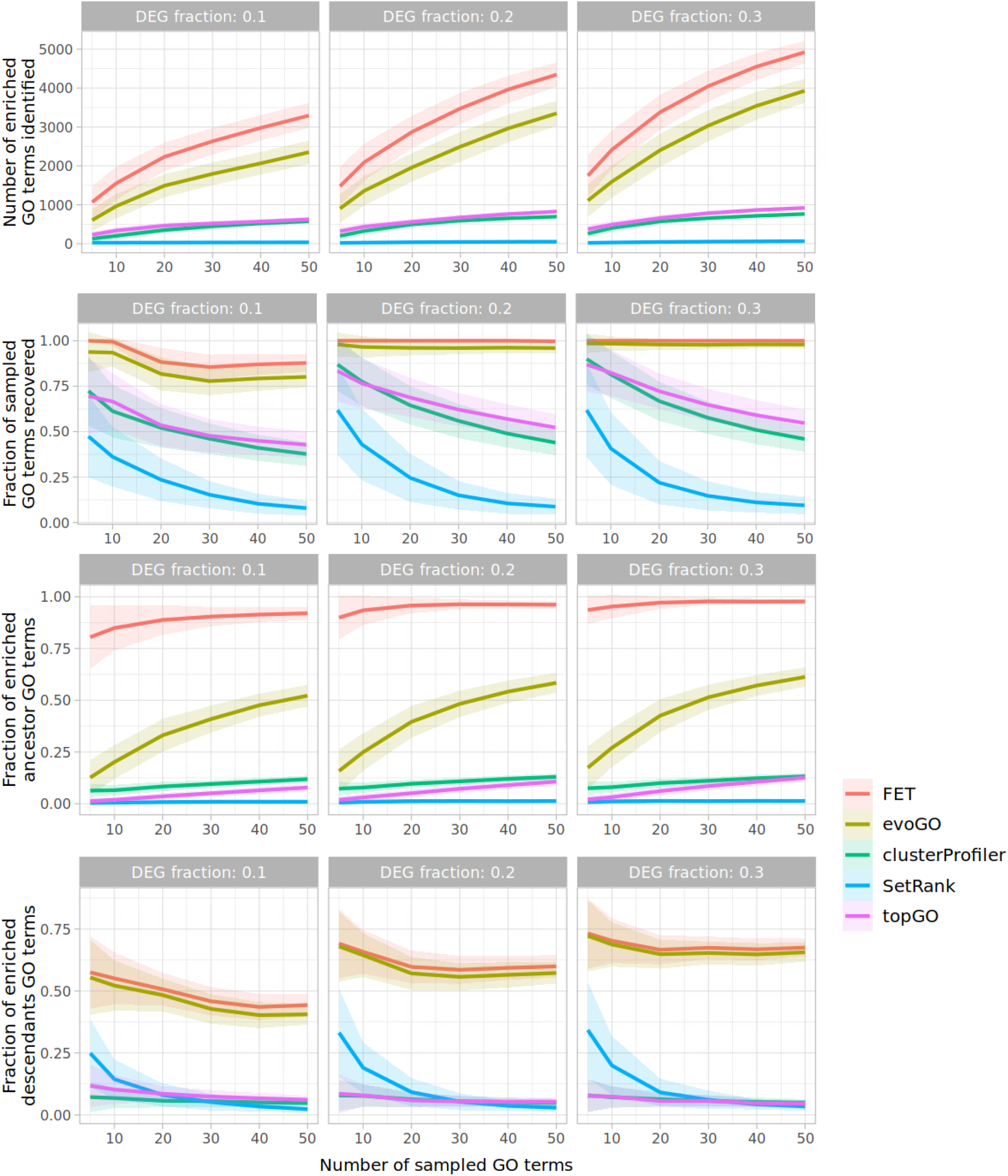
Results of synthetic benchmark. **(A)** Total number of significantly enriched GO terms per GO enrichment analysis method; **(B)** fraction of significantly enriched randomly sampled (true positive) GO terms; **(C, D)** fraction of enriched ancestors and descendants of the sampled terms among all ancestors and descendants, correspondingly. The lines and bands indicate mean±SD values per method. GO terms were randomly sampled 500 times.

topGO, clusterProfiler, and SetRank yielded much shorter lists of enriched terms than FET (by 79, 83 and 99%, correspondingly). However, these methods also recovered significantly fewer TP terms (65, 61 and 26%, correspondingly). In contrast, on average, evoGO recovered 96% of TPs detected by FET, while the total number of significantly enriched terms was reduced by 30%.

All the tested advanced methods significantly reduced the number of enriched ancestors of TP terms compared to FET. This illustrates that to a certain extent, all of these methods decreased the redundancy of the enrichment analysis results. topGO, clusterProfiler, and SetRank produced similar results and reduced the number of enriched ancestors by more than 85%. On the other hand, the down-weighting of ancestor GO terms by evoGO was less aggressive (by up to 57%) and more dependent on the total number of DE genes. evoGO also did not affect the number of enriched descendant terms, which was expected as the method only modifies the ancestor gene weights. Other methods, despite being based on fundamentally different principles, strongly suppressed descendants of TP GO terms (Fig. 1).

Considering our findings, we investigated the extent to which the outcome of the benchmark might be influenced by the selected p-value threshold. It is possible that higher threshold values may result in a higher fraction of recovered TPs for some methods. To test this, we reran the experiment with a fixed number of sampled terms, but using a variable significance threshold. Altering the p-value threshold across a wide range (0.01-0.2) does not significantly affect the outcome of the synthetic benchmark (Supplementary Fig. 1). Even with higher p-value thresholds, topGO, SetRank, and clusterProfiler do not reach the TP rate achieved by evoGO.

In summary, in our synthetic benchmark, evoGO demonstrates a less aggressive approach to reducing redundancy compared to other methods, but retains a notably higher percentage of recovered true positives, regardless of the p-value threshold.

## Real data benchmark

The benchmark using real data shows trends consistent with those observed in the synthetic test. FET returned the most extensive list of enriched GO terms and also identified the highest number of TS terms. evoGO demonstrated the second-highest fraction of recovered TS GO terms, whereas the number of enriched terms identified by evoGO was reduced by approximately 30-40% compared to FET. In line with the synthetic benchmark, the remaining methods detected significantly fewer enriched terms and recovered a much lower number of terms of interest. Despite this, the average proportion of true positive and false positive TS terms in the enriched terms was roughly equivalent across all methods (Fig. 2).

**Fig. 2.**
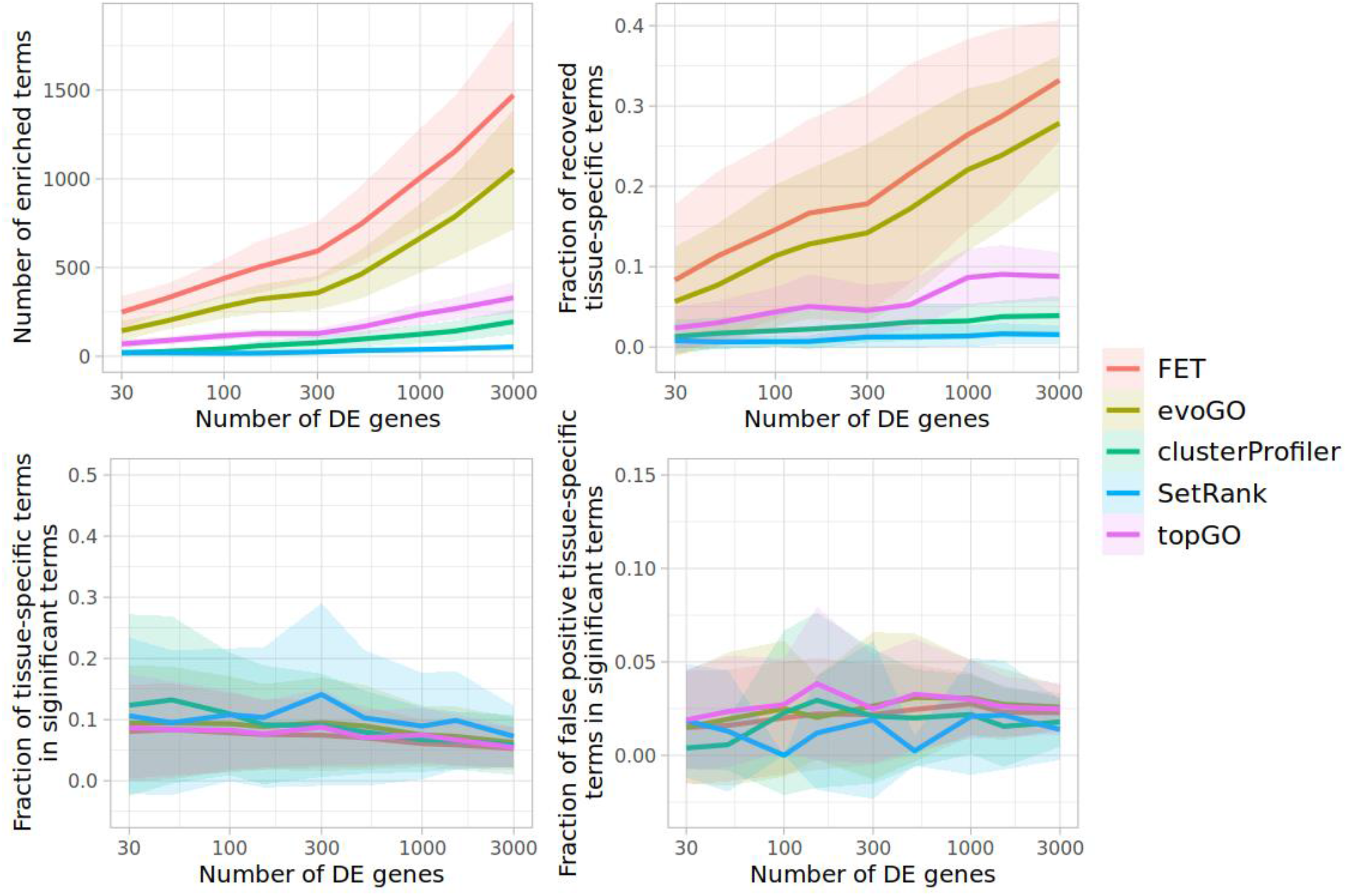
Results of real data benchmark. **(A)** Total number of significantly enriched GO terms per GO enrichment analysis method; **(B)** fractions of significantly enriched TS GO terms; **(C, D)** fractions of true-positive and false-positive TS terms in all significantly enriched GO terms. The lines and bands indicate mean±SD values per method. The values were obtained using comparisons of gene expression profiles of healthy human tissues (see Methods).

To ensure that our choice of p-value threshold did not influence the outcome of the benchmark, we validated the results using variable p-value threshold. We found that irrespective of the threshold set, the proportion of recovered TS terms for FET and evoGO is significantly higher than that of other methods (Supplementary Fig. 2).

While the number of recovered TS terms is an important metric, it alone does not provide a comprehensive measure of a GO enrichment method’s performance under the given benchmark conditions. All the tested advanced methods aim to reduce the number of redundant enriched GO terms in the analysis results. Therefore, a lower number of terms might provide a better representation of the involved biological processes. In view of this, we also evaluated the fractions of ancestors of TS GO terms that did not reach significance. We assumed that TS terms are among the key terms to identify when comparing gene expression profiles of different tissues. Thus, they should only be deprioritized in favor of other more informative (descendant) TS terms. Subsequently, if some of the terms were not found among significant terms and their non-significant ancestors, it could indicate that they were eliminated without adequate justification.

Considering that all tested methods are based on FET, we used the set of significant TS terms and their ancestors recovered by this method as a reference.

The results in Fig. 3 demonstrate that evoGO effectively reduces the number of identified TS terms by down-weighting the ancestors of other TS terms. This is expected, as higher-level terms are typically considered redundant. In contrast, other advanced GO enrichment analysis methods eliminate a significant number of TS terms, even if they lack significant TS descendants. As previously mentioned, we believe this could lead to the loss of valuable information during analysis.

**Fig. 3.**
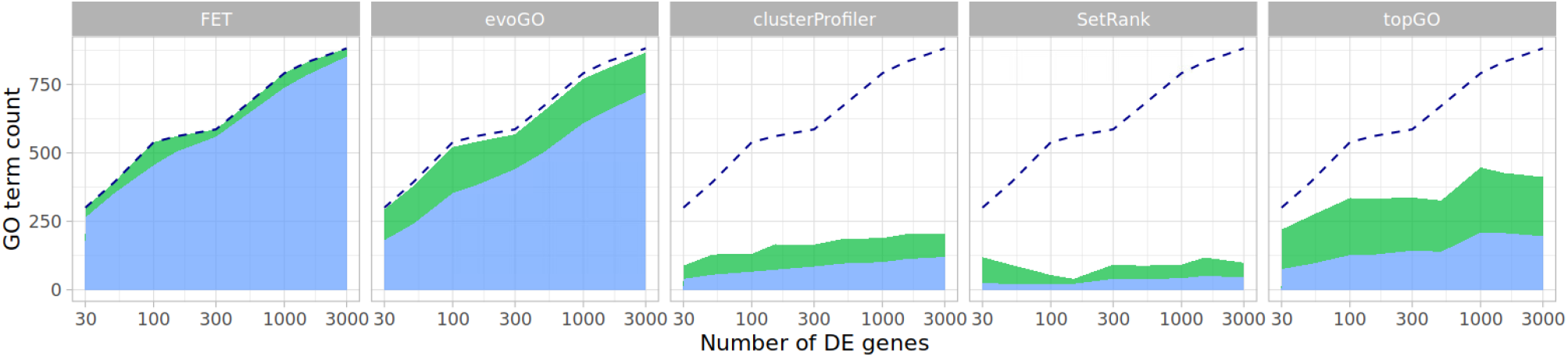
Recovered TS GO terms and their non-recovered ancestors. The number of recovered TS terms is represented by the blue area, and the number of non-recovered ancestors is represented by the green area. Considering that all tested methods are based on FET, a dashed line indicating the sums of these values, as identified by FET, is shown in each plot for comparison.

Following the same logic, we also counted significant terms that had no significant ancestors or descendants but were eliminated by the tested methods. Consistent with the results described above, evoGO shows the lowest number of missing “orphan” terms, closely followed by topGO. In contrast, SetRank and clusterProfiler show significantly more aggressive GO term down-weighting (Fig. 4).

**Fig. 4.**
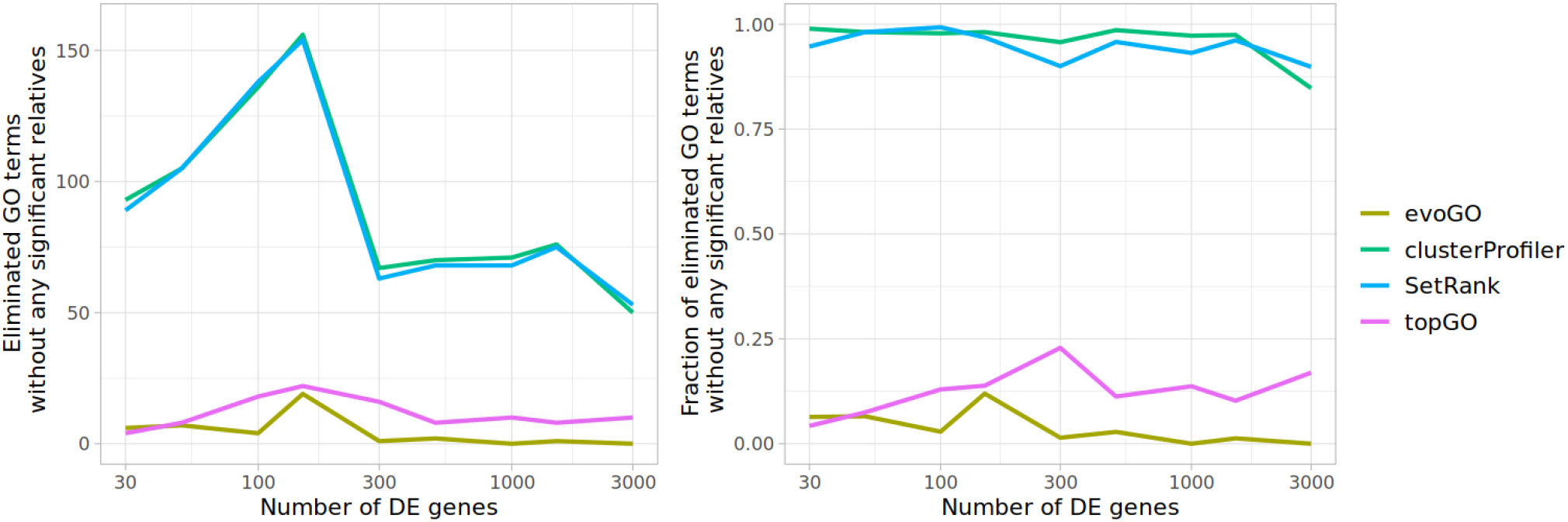
Deprioritization of significantly enriched terms without significant relatives. **(A)** Total number of significant terms, which had no significant ancestors or descendants, but were deprioritized and lost their significance as compared to the results of FET; **(B)** same as (A), but the data are shown as a fraction of the total number of significant terms without enriched relatives, as identified by FET.

Finally, considering that all tested methods eliminated both TS and non-TS GO terms to some extent, we evaluated whether the methods showed any preference for preserving TS terms. Specifically, we calculated the ratio between the fraction of eliminated TS and non-TS terms. We find this ratio to be a more representative measure of the methods’ ability to prioritize biologically relevant terms than, for example, precision, due to the very low ratio of TS to non-TS enriched terms. As can be observed in Fig. 5, in all cases, the ratio was lower than 1, suggesting that all tested methods indeed demonstrated certain selectivity towards TS terms. Furthermore, evoGO’s ratio was almost 2 times lower compared to the other methods. Overall, this finding suggests that evoGO was more efficient in prioritizing biologically relevant GO terms.

**Fig. 5.**
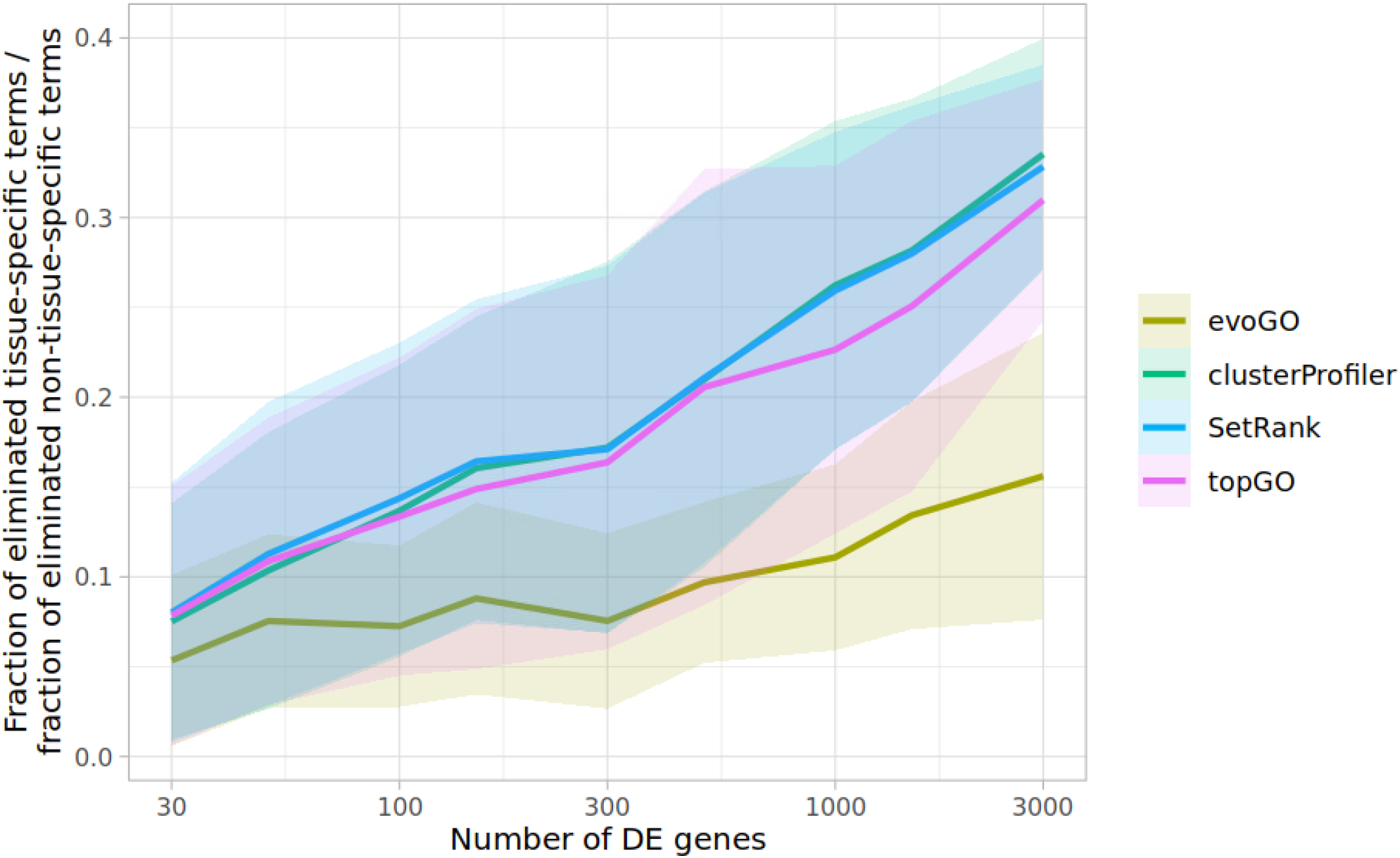
GO term elimination selectivity. The values represent the ratio between the fractions of eliminated TS and non-TS terms for each tested method. Terms were considered eliminated if they lost their significance as compared to the results obtained with FET. The lines and bands indicate the mean±SD values per method.

### Execution time comparison

Short analysis execution times are crucial for large-scale projects such as compound screening, where hundreds or even thousands of GO enrichment analyses need to be performed within a reasonable timeframe. Therefore, we also compared the time taken to perform an analysis for each method, utilizing both synthetic and real-life data. In general, we found significant differences in execution times depending on the analysis method and the number of DE genes (Fig. 6). As expected, FET was the fastest method due to its simplicity. On the other hand, topGO was one of the slowest, taking more than 3 minutes per analysis in most scenarios. On average, the evoGO analysis took 37 seconds with real data and 52 seconds with synthetic data. Notably, the execution times of SetRank and clusterProfiler were significantly longer in the synthetic benchmark than in the real data benchmark (up to 5.8 min for SetRank and 15 min for clusterProfiler). Apparently, these methods are sensitive to the distribution of DE genes across GO terms. In addition, the speed of SetRank strongly depended on the selected maximum gene set size considered during the analysis. In our analyses, this value was set to 300, which is the default chosen by the authors. However, increasing the maximum size of gene sets to accommodate all GO terms can significantly prolong the analysis duration (up to 45 minutes).

**Fig. 6.**
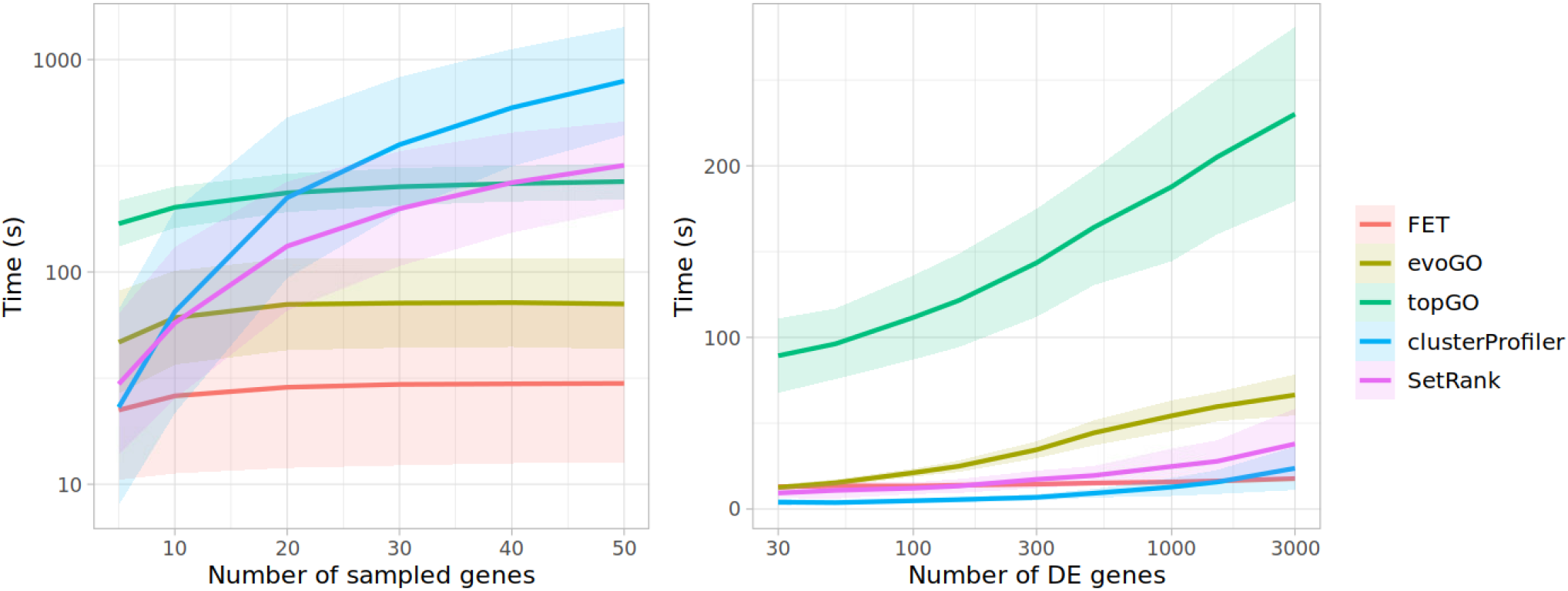
Comparison of GO enrichment analysis execution time. **(A)** Analysis execution time depending on the number of randomly sampled GO terms in the synthetic benchmark; **(B)** Analysis execution time depending on the number of DE genes in the real data benchmark. The lines and bands indicate mean±SD values per method.

## Conclusions

Before drawing the conclusions, we would like to mention certain limitations of this study. First of all, the synthetic benchmark does not allow us to estimate the false positive rates based on the results provided by the tested methods. This stems from the fact that TP GO terms were selected randomly and without a biological ground truth, it is impossible to classify other terms sharing the same genes as “true positives” or “false positives”. On the other hand, in the real data benchmark, the GO terms associated with irrelevant tissues were considered false positives. However, the differences between methods in the recovery of these false positive terms were not conclusive due to the high variability of results.

In the real data benchmark, it might be seen as somewhat subjective to assume that TS GO terms are the key terms to be discovered when comparing gene expression across different tissues. It is plausible that some non-TS terms also correctly reflect the differences between tissues, and thus, their deprioritization could potentially overlook important biological insights. Nevertheless, we expect that TS GO terms are still relevant and their correct identification is a good indicator of a method’s performance.

In our benchmarks, we evaluated the ability of different GO enrichment analysis methods to de-prioritize enriched GO terms that may be considered redundant. All tested methods, despite being based on the same FET, showed dramatically different performance in terms of accuracy and analysis execution times. topGO, clusterProfiler, and SetRank significantly reduced the number of GO terms that are identified by FET as significantly enriched.

However, this strongly affected their ability to recover true positive terms in both synthetic and real data benchmark. SetRank showed the most aggressive term elimination.

Typically, it recovered less than 10% of terms found to be significantly enriched by FET. In contrast, evoGO’s redundant term elimination was the least destructive and the most selective among the tested GO enrichment analysis methods. In particular, evoGO most

efficiently prioritized TS GO terms in the analyses, which were expected to capture the biological processes inherent to various human tissues. Furthermore, in those analyses, evoGO eliminated the fewest enriched GO terms that had no significantly enriched relatives and thus should not normally be considered redundant. This demonstrates that evoGO has the best potential for preserving valuable information among all the tested methods.

In addition to these findings, we would like to note that p-values provided by evoGO are more easily interpretable than those provided by methods like topGO or SetRank. evoGO calculates the initial p-values using FET and they stay unchanged for the most significant GO terms in their ancestries. As a result, p-values for non-down-weighted terms preserve their meaning, and the standard p-value thresholds should still be applicable for evoGO results. This makes filtering for significance both straightforward and consistent with conventional statistical practices.

evoGO showed the shortest execution times among the advanced methods in the synthetic benchmark and outperformed the other GO hierarchy-based method, topGO, in the real data benchmark. Even though evoGO was not the fastest method in all scenarios, its execution times remained consistent in both benchmarks, which was not the case for other methods. Based on our experience, evoGO’s speed is sufficient for performing large-scale studies comprising thousands of analyses within a reasonable timeframe.

In conclusion, our findings demonstrate that among FET-based approaches evoGO stands out as an effective and fast method for minimizing redundancy in GO enrichment analysis results while reliably preserving biologically relevant information. We believe that evoGO could be a valuable tool for conducting downstream analyses, particularly in the context of high-throughput transcriptomic and proteomic screening studies.

## REFERENCES

1. Minoru Kanehisa, Miho Furumichi, Mao Tanabe, Yoko Sato, Kanae Morishima, KEGG: new perspectives on genomes, pathways, diseases and drugs, Nucleic Acids Research, Volume 45, Issue D1, January 2017, Pages D353–D361, 10.1093/nar/gkw1092

2. Antonio Fabregat, Steven Jupe, Lisa Matthews, Konstantinos Sidiropoulos, Marc Gillespie, Phani Garapati, Robin Haw, Bijay Jassal, Florian Korninger, Bruce May, Marija Milacic, Corina Duenas Roca, Karen Rothfels, Cristoffer Sevilla, Veronica Shamovsky, Solomon Shorser, Thawfeek Varusai, Guilherme Viteri, Joel Weiser, Guanming Wu, Lincoln Stein, Henning Hermjakob, Peter D’Eustachio, The Reactome Pathway Knowledgebase, Nucleic Acids Research, Volume 46, Issue D1, 4 January 2018, Pages D649–D655, 10.1093/nar/gkx1132

3. Arthur Liberzon, Chet Birger, Helga Thorvaldsdóttir, Mahmoud Ghandi, Jill P. Mesirov, Pablo Tamayo. The Molecular Signatures Database Hallmark Gene Set Collection, Cell Systems, Volume 1, Issue 6, 2015, Pages 417–425, 10.1016/j.cels.2015.12.004

4. The Gene Ontology Consortium, The Gene Ontology Resource: 20 years and still GOing strong, Nucleic Acids Research, Volume 47, Issue D1, 08 January 2019, Pages D330–D338, 10.1093/nar/gky1055

5. Bayerlová, M., Jung, K., Kramer, F. et al. Comparative study on gene set and pathway topology-based enrichment methods. BMC Bioinformatics 16, 334 (2015). 10.1186/s12859-015-0751-5

6. Joanna Zyla, Michal Marczyk, Teresa Domaszewska, Stefan H E Kaufmann, Joanna Polanska, January Weiner, Gene set enrichment for reproducible science: comparison of CERNO and eight other algorithms, Bioinformatics, Volume 35, Issue 24, December 2019, Pages 5146–5154, 10.1093/bioinformatics/btz447

7. Ludwig Geistlinger, Gergely Csaba, Mara Santarelli, Marcel Ramos, Lucas Schiffer, Nitesh Turaga, Charity Law, Sean Davis, Vincent Carey, Martin Morgan, Ralf Zimmer, Levi Waldron, Toward a gold standard for benchmarking gene set enrichment analysis, Briefings in Bioinformatics, Volume 22, Issue 1, January 2021, Pages 545–556, 10.1093/bib/bbz158

8. Bioconductor’s EnrichmentBrowser: seamless navigation through combined results of set-& network-based enrichment analysis. 10.1186%2Fs12859-016-0884-1

9. Irizarry RA, Chi Wang, Yun Zhou, Speed TP. Gene set enrichment analysis made simple. Statistical Methods in Medical Research. 2009;18(6):565–575. doi:10.1177/0962280209351908

10. Simillion, C., Liechti, R., Lischer, H.E. et al. Avoiding the pitfalls of gene set enrichment analysis with SetRank. BMC Bioinformatics 18, 151 (2017). 10.1186/s12859-017-1571-6

11. Wang, G., Oh, DH. & Dassanayake, M. GOMCL: a toolkit to cluster, evaluate, and extract non-redundant associations of Gene Ontology-based functions. BMC Bioinformatics 21, 139 (2020). 10.1186/s12859-020-3447-4

12. Jantzen, S.G., Sutherland, B.J., Minkley, D.R. et al. GO Trimming: Systematically reducing redundancy in large Gene Ontology datasets. BMC Res Notes 4, 267 (2011). 10.1186/1756-0500-4-267

13. Frida Belinky, Noam Nativ, Gil Stelzer, Shahar Zimmerman, Tsippi Iny Stein, Marilyn Safran, Doron Lancet, PathCards: multi-source consolidation of human biological pathways, Database, Volume 2015, 2015, bav006, 10.1093/database/bav006

14. Juan C. Vivar, Priscilla Pemu, Ruth McPherson, and Sujoy Ghosh, Redundancy Control in Pathway Databases (ReCiPa): An Application for Improving Gene-Set Enrichment Analysis in Omics Studies and “Big Data” Biology, OMICS: A Journal of Integrative Biology, Vol. 17, No. 8, 2013. 10.1089%2Fomi.2012.0083

15. Stoney, R.A., Schwartz, JM., Robertson, D.L. et al. Using set theory to reduce redundancy in pathway sets. BMC Bioinformatics 19, 386 (2018). 10.1186/s12859-018-2355-3

16. Sara R. Savage, Zhiao Shi, Yuxing Liao, Bing Zhang, Graph Algorithms for Condensing and Consolidating Gene Set Analysis Results, Molecular & Cellular Proteomics, Volume 18, Issue 8, Supplement 1, 2019, Pages S141–S152, 10.1074/mcp.TIR118.001263.

17. Yu, G. (2020). Gene Ontology Semantic Similarity Analysis Using GOSemSim. In: Kidder, B. (eds) Stem Cell Transcriptional Networks. Methods in Molecular Biology, vol 2117. Humana, New York, NY. 10.1007/978-1-0716-0301-7_11

18. Zuguang Gu, Daniel Hübschmann, SimplifyEnrichment: A Bioconductor Package for Clustering and Visualizing Functional Enrichment Results, Genomics, Proteomics & Bioinformatics, Volume 21, Issue 1, February 2023, Pages 190–202, 10.1016/j.gpb.2022.04.008

19. Adrian Alexa, Jörg Rahnenführer, Thomas Lengauer, Improved scoring of functional groups from gene expression data by decorrelating GO graph structure, Bioinformatics, Volume 22, Issue 13, July 2006, Pages 1600–1607, 10.1093/bioinformatics/btl140

20. Tarca, A.L., Draghici, S., Bhatti, G. et al. Down-weighting overlapping genes improves gene set analysis. BMC Bioinformatics 13, 136 (2012). 10.1186/1471-2105-13-136

21. Tianzhi Wu, Erqiang Hu, Shuangbin Xu, Meijun Chen, Pingfan Guo, Zehan Dai, Tingze Feng, Lang Zhou, Wenli Tang, Li Zhan, Xiaocong Fu, Shanshan Liu, Xiaochen Bo, Guangchuang Yu, clusterProfiler 4.0: A universal enrichment tool for interpreting omics data, The Innovation, Volume 2, Issue 3, 2021, 10.1016/j.xinn.2021.100141.

22. Lonsdale, J., Thomas, J., Salvatore, M. et al. The Genotype-Tissue Expression (GTEx) project. Nat Genet 45, 580–585 (2013). 10.1038/ng.2653

23. Love, M.I., Huber, W. & Anders, S. Moderated estimation of fold change and dispersion for RNA-seq data with DESeq2. Genome Biol 15, 550 (2014). 10.1186/s13059-014-0550-8

