## Supplementary for "Cutting through the clutter: minimizing redundancy in GO enrichment analysis with evoGO"

### SUPPLEMENTARY MATERIALS

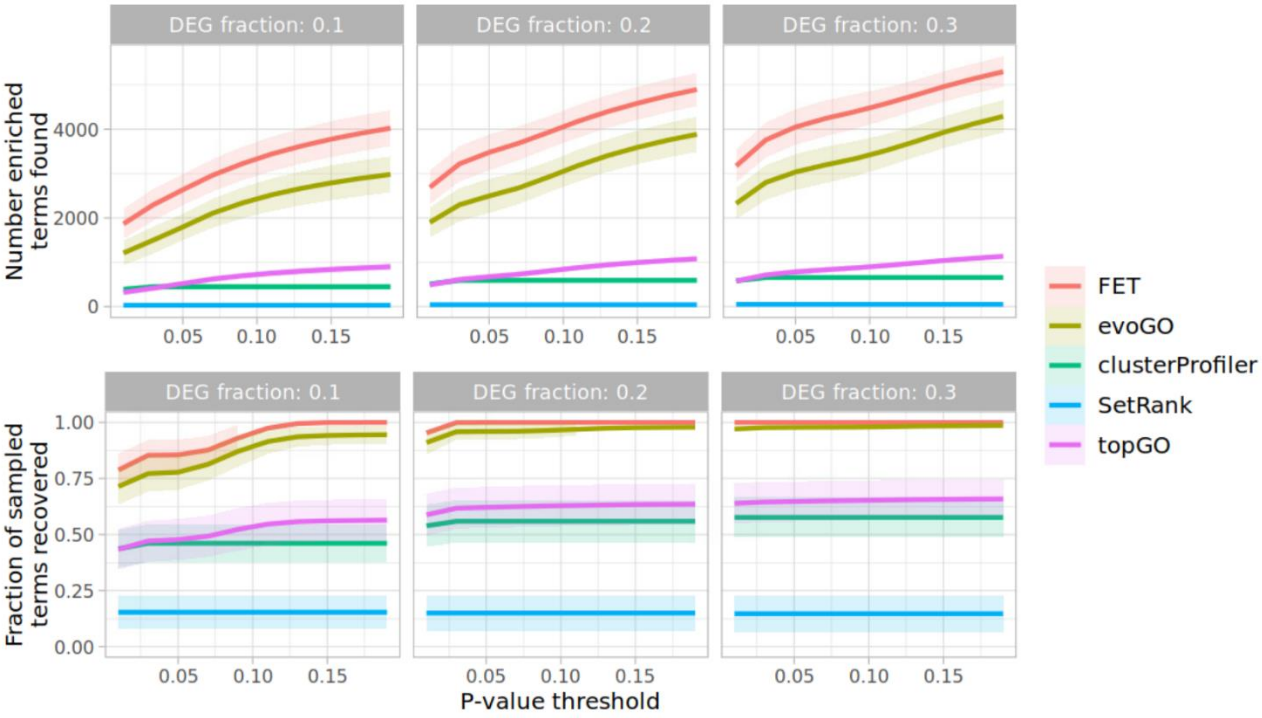


**Supplementary Fig. 1. The effect of varying p-value threshold on the outcome of the synthetic benchmark. (A)** Total number of significantly enriched GO terms; **(B)** fraction of significantly enriched randomly sampled (true positive) GO terms. The lines and bands indicate mean±SD values per method. The results were obtained with 30 sampled GO terms. GO terms were randomly sampled 500 times.

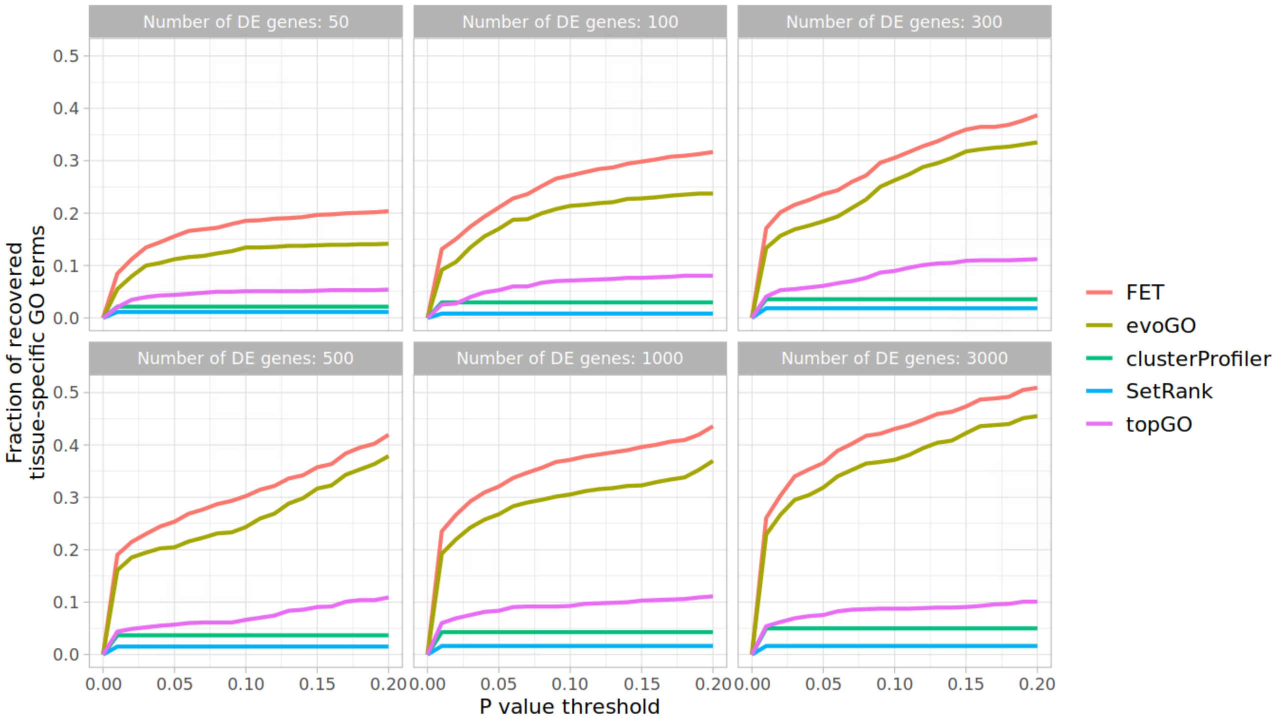


**Supplementary Fig. 2. The effect of varying the p-value threshold on the fraction of recovered tissue-specific GO terms.** The values represent fractions of the total number of tissue-specific terms available for tissues that were used in the benchmark.
